# Brain structural modules associated to functional high-order interactions in the human brain

**DOI:** 10.1101/2025.03.21.644509

**Authors:** Borja Camino-Pontes, Antonio Jimenez-Marin, Iñigo Tellaetxe-Elorriaga, Izaro Fernandez-Iriondo, Asier Erramuzpe, Ibai Diez, Paolo Bonifazi, Marilyn Gatica, Fernando E. Rosas, Daniele Marinazzo, Sebastiano Stramaglia, Jesus M. Cortes

## Abstract

The brain’s modular organization, ranging from microcircuits to large-scale networks, has been extensively studied in terms of its structural and functional properties. Particularly insightful has been the investigation of the coupling between structural connectivity (SC) and functional connectivity (FC), whose analysis has revealed important insights into the brain’s efficiency and adaptability related to various cognitive functions. Interestingly, links in SC are intrinsically pairwise but this is not the case for FC; and while recent work demonstrates the relevance of the brain’s high-order interactions (HOI), the coupling of between SC and functional HOI remains unexplored. To address this gap, this study leverages functional MRI and diffusion weighted imaging to delineate the brain’s modular structure by investigating the coupling between SC and functional HOI. Our results demonstrates that structural networks can be associated with both redundant and synergistic functional interactions. In particular, SC exhibits both positive and negative correlations with redundancy, it shows consistent positive correlations with synergy, indicating that a higher density of structural connections is linked to increased synergistic interactions. These findings advance our understanding of the complex relationship between structural and high-order functional properties, shedding light on the brain’s architecture underlying its modular organization.

**Significance Statement:** Understanding the intricate relationship between the brain structural and functional connectivity is instrumental for enabling a richer characterization of cognitive functions. This challenge is complex, since it involves comparing pairwise links representing white-matter fibers with higher-order functional interactions that can involve multiple brain regions sharing information — indeed, these interactions can be redundant in presence of shared information, or synergistic when novel information arises in the whole subsystem. Here we advance the investigation of the coupling between brain structure and high-order interactions, offering new insights into how structural properties are associated with both redundant and synergistic functional modules. Furthermore, the approach presented here opens new doors to investigate how this coupling between structural connectivity and functional HOI is modulated during development or under diverse brain pathologies.

## Introduction

The brain modular organization has been extensively studied across different scales, from local microcircuits to large-scale brain networks (Bassett and E. T. Bullmore 2009; P. Bonifazi et al. 2009; E. Bullmore and Sporns 2009; Meunier 2009; Zalesky et al. 2010; Feldt et al. 2011; Sporns 2011; Craddock et al. 2012; Martín-Suárez et al. 2023; Clusella et al. 2023). These investigations show that brain modules are not isolated but instead interact with each other to facilitate information processing and long-distance communication within the brain. These complex interactions are reflected in the brain’s structure and function, and can be investigated via the structural connectivity (SC) and functional connectivity (FC) (Koch et al. 2002; Honey et al. 2007; Crofts and Higham 2009; Deco et al. 2009; Park and Friston 2013; Suárez et al. 2020).

The study of structure-function coupling (SFC), understood as associations between the brain regions’ functional activity and the corresponding white-matter architecture, have provided important insights into the remarkable efficiency and adaptability of the brain (see (Fotiadis, Parkes, et al. 2024) and references therein). Extending conventional approaches that examine link-to-link relationships between SC and FC, several studies have focused on modular coupling reflecting the interactions between SC and FC at the level of modules or communities (Diez et al. 2015; Sporns and Betzel 2016; Betzel et al. 2019; Fernandez-Iriondo et al. 2021; Puxeddu et al. 2022; Jimenez-Marin et al. 2024). Interestingly, while links in SC are intrinsically pairwise (due to white-matter fibers projecting from one region to another), this is not necessarily the case in FC. Yet, most FC analyses focus on pairwise statistics and neglect higher order interactions (HOI), whose relevance is increasingly being recognized in relation to healthy aging (Camino-Pontes et al. 2018; Gatica, Cofré, et al. 2021; Gatica, E. Rosas, et al. 2022), higher-level cognitive tasks (Andrea I. Luppi, P. A. M. Mediano, et al. 2022), epilepsy (Erramuzpe et al. 2015), neurodegeneration (Herzog et al. 2022), and the functioning of anesthesia (Andrea I Luppi, P. A. Mediano, et al. 2024; Andrea I. Luppi, Fernando E. Rosas, et al. 2024b). While results have shown that the activity of two regions (including lagged interactions) tends to become redundant when they are structurally connected (Andrea I. Luppi, P. A. M. Mediano, et al. 2022; Gatica, Atkinson-Clement, et al. 2024), a systematic investigation of the relationship between high-order FC and SC at a modular level is still missing.

To fill this gap, here we investigate the relation between SC and functional HOI. We calculate functional HOI via the O-information (Fernando E. Rosas et al. 2019; Stramaglia et al. 2021), which can quantify the predominance of redundancy or synergy in modules of three or more brain regions — the former corresponding to when information is stored in multiple regions, and the latter when information is not stored in any specific part but rather in the whole (Andrea I Luppi, Fernando E Rosas, et al. 2024a). We complement the global assessment provided by the O-information by also studying its gradients, which have been recently introduced as low-order descriptors that characterize the role of individual regions in high-order effects (Scagliarini, Nuzzi, et al. 2023).

## Methodology

### Experimental Design

#### Neuroimaging Dataset

Neuroimaging data was shared by the Max Planck Institute Leipzig Mind-Brain-Body Dataset, under the name of LEMON (Babayan et al. 2018; Babayan et al. 2019), a rich multimodal dataset comprising MRI sequences, EEG, ECG, and behavioral scores. For this study, we focused on a subset of N = 136 healthy individuals aged between 20 and 30 years, with males constituting 98 individuals (72%). The selection of this group of subjects was done to avoid known age-effects in both structural and functional connectivity (Paolo Bonifazi et al. 2018). Specifically, we have used here three MRI sequences: T1, DWI and rs-fMRI. For detailed descriptions of these sequences, please refer to Table 7 in (Babayan et al. 2019).

#### Brain Imaging Preprocessing

The functional HOI were assessed from time-series data provided by LEMON. Although the preprocessing details can be found in (Mendes et al. 2019), major steps were: 3D motion correction (FSL MCFLIRT) (Jenkinson, Bannister, et al. 2002), distortion correction (FSL FUGUE) (Jenkinson, Beckmann, et al. 2012), rigid-body co-registration of the unwrapped temporal mean image to the individual’s anatomical image (FreeSurfer bbregister) (Greve and Fischl 2009), denoising (Nipype rapidart and aCompCor) (Behzadi et al. 2007), band-pass filtering between 0.01-0.1 Hz (FSL), mean centering, as well as variance normalization of the denoised time series (Nitime) (Rokem et al. 2009), and spatial normalization to MNI152 at 2 mm resolution via transformation parameters derived during structural preprocessing (ANTs SyN) (Avants et al. 2011). We further used two tools to address physiological effects such as systemic low frequency oscillations (sLFOs) and the hemodynamic response function. Rapidtide (Blaise deB et al. 2024) was used to detect lagged signal propagation patterns within the BOLD signal, correcting for inflated spurious correlations in the data (Korponay et al. 2024). The hemodynamic response function at rest was estimated with a blind procedure based on point process and a mixture of Gamma functions as informed basis set (Wu et al. 2021) and deconvolved, eliminating temporal confounds (Rangaprakash et al. 2018).

The SC obtained from DWI data was preprocessed following the detailed procedure reported in (Jimenez-Marin et al. 2024) and publicly available in (Jimenez-Marin 2024), consisting of cleaning the raw images using *DWIdenoise* and *DWIpreproc* tools from MRtrix3 (J-Donald Tournier et al. 2019), which included correction for eddy current distortion and susceptibility-induced distortion using the *topup* tool from FSL, whole-brain tractography by the *iFOD2* tracking algorithm in MRtrix3 (Jacques-Donald Tournier et al. 2010). The tractography was initiated from seeds located at the grey matter-white matter mask, with a selection of 3 million fibers based on an angle threshold of *<*45º and a length threshold of *<*200mm, following the MRtrix3 documentation. Finally, we also corrected the effect of crossing and disconnection of fibers in the quantification of the total number of streamlines using the tool *SIFT2* (Smith et al. 2015). This process provided SC for each subject, which represents pairwise connectivity strength, calculated as the sum of streamline weights connecting regions pairs.

#### Brain Modules

Brain networks need a definition for nodes and links. In this study, we used the Desikan-Killiany atlas (Desikan et al. 2006), which comprises 68 cortical regions (34 per hemisphere), to define network nodes. These 68 regions were further grouped into four modules (or macroregions) based on anatomical lobes: Temporal (TEM), Frontal (FRO), Occipital (OCC), and Parietal (PAR), facilitating modular analyses. For HOI analyses, we extracted a representative time series for each node by averaging the time series of all voxels within that region. The same node definitions were used to construct the SC matrices.

As a control, to verify that some of our analyses did not depend on the specific choice of modular structure, we also employed an alternative functional atlas. Specifically, we used the Schaefer atlas (Schaefer et al. 2018), which consists of 100 nodes (50 in the right hemisphere and 50 in the left hemisphere). These 100 nodes were grouped into larger macroregions for modular analysis. For this study, we selected the 7 Yeo resting-state networks Thomas Yeo et al. 2011: visual (VIS), somatomotor (SM), dorsal attention (DA), ventral attention (VA), limbic (LIM), frontoparietal (FP), and default mode networks (DMN).

To extend both the functional and anatomical atlases beyond cortical regions, we incorporated additional subcortical (SUB) macroregions using the FreeSurfer parcellation Fischl 2012. This integration introduced 16 subcortical nodes, with 8 nodes in the right hemisphere and 8 in the left hemisphere. The selected subcortical regions included the thalamus (THA), caudate (CAU), putamen (PUT), pallidum (PAL), hippocampus (HIP), amygdala (AMY), accumbens (ACC), and ventral diencephalon (vDI). Additionally, two cerebellar nodes, one for the left hemisphere and one for the right, were incorporated into both the functional and anatomical atlases.

### Statistical Analysis

#### Measuring functional high-order interactions via the O-information

The activity of the time series data from different brain regions is represented as random variables *X*_1_, *X*_2_, etc. The average information contained in those variables can be estimated via Shannon’s entropy *H*(*X*) = − ∑_*x*_ prob(*x*)log(prob(*x*)), where *x* represents the possible states of variable *X* (Cover and Thomas 2005). The part of that information that is shared between pairs of variables *X*_1_ and *X*_2_ is measured by the mutual information *I*(*X*_1_, *X*_2_) = *H*(*X*_1_, *X*_2_) − *H*(*X*_1_) − *H*(*X*_2_), where *H*(*X*_1_, *X*_2_) is the joint entropy. For three variables, the difference between pairwise and higher-order effects can be measured via the interaction information *II*(*X*_1_; *X*_2_; *X*_3_) = *I*(*X*_1_; *X*_3_) + *I*(*X*_2_; *X*_3_) − *I*(*X*_1_, *X*_2_; *X*_3_) (McGill 1954), where *I*(*X*_1_, *X*_2_; *X*_3_) is the mutual information between (*X*_1_, *X*_2_) and *X*_3_. Interestingly, *II*(*X*_1_; *X*_2_; *X*_3_) is symmetric in its three variables (Erramuzpe et al. 2015). For interactions involving more than three variables, the balance between low and high order effects can be captured via the O-information (Fernando E. Rosas et al. 2019)

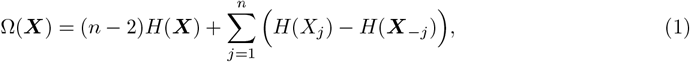

where ***X*** ≡ (*X*_1_, *X*_2_, …, *X*_*N*_) are the *n* random variables and ***X***_−*j*_ ≡ (*X*_1_, …, *X*_*j*−1_, *X*_*j*+1_, …, *X*_*N*_) is equal to ***X*** but removing the component *X*_*j*_. Note that for *n* = 3 the O-information coincides with the interaction information. The sign of the O-information determines whether the interactions in the group ***X*** are dominated by redundancy (Ω(***X***) *>* 0), or if they are dominated by synergy (Ω(***X***) *<* 0).

Our analyses focus on modules of *n* = 3. For each subject and triplet (*X*_1_, *X*_2_, *X*_3_), we calculate the O-information by estimating the entropy using a rank-based normalization allowing a parsimonious estimate suitable for Gaussian processes (Ince et al. 2017). To assess the statistical significance of the computed O-information values, we employed a bootstrapping approach with 1000 surrogates. For each subject and triplet, we generated shuffled datasets and recomputed the O-information, allowing us to construct confidence intervals and evaluate deviations from a null distribution. The code to compute the O-information is publicly available at (Marinazzo 2024; Combrisson 2025).

#### Gradients of O-information

We complement the global assessment provided by the O-information by additional analyses considering the gradients of the O-information (Scagliarini, Nuzzi, et al. 2023), which determine the contribution of individual variables (or pairs, triplet, etc) to a specific pattern of HOI. Specifically, the gradient for a triplet of variables (*X*_*i*_, *X*_*j*_, *X*_*k*_) is given by

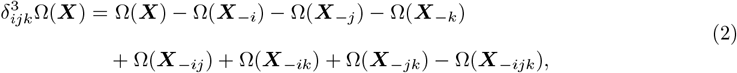

where ***X***_−*ij*_ denotes ***X*** after removing the variables *X*_*i*_ and *X*_*j*_, and ***X***_−*ijk*_ denotes ***X*** after removing the triplet of variables *X*_*i*_, *X*_*j*_ and *X*_*k*_. The gradients of O-information allowed us to evaluate the contribution of a specific triplet (*X*_*i*_, *X*_*j*_, *X*_*k*_) to the entire system, a contribution that cannot be attributed to the inclusion of pairs or individual variables within the triplet. The sign of the gradient can be interpreted similarly as in the O-information, with 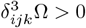 implying a redundancy-dominated contribution of the triplet, and 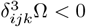 for a synergy-dominated one. The gradients of O-information were calculated for each subject and triplet using the publicly available code at (Marinazzo 2022).

#### SC-Functional HOI Coupling

To investigate the modular coupling between SC and functional HOI, we defined modules by using the different macroregions in the brain atlases described before. For each subject, we calculated the functional HOI for all possible triplets of node time series, obtaining a 3D matrix (tensor), where its elements correspond with the O-information values for a specific triplet. For both SC and HOI matrices, we first calculated population representation by considering for each element in the matrix the median value across individuals in the population. Following previous work (Preti and Van De Ville 2019; Liégeois et al. 2020; Gu et al. 2021; Fotiadis, Cieslak, et al. 2023), we defined the coupling as the Spearman correlation of the structure elements within the module 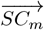, equal to the vector-wise SC between pairs of nodes (*X*_*i*_, *X*_*j*_) in the macroregion *m*) and HOI elements represented by 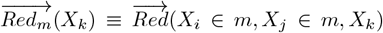 and 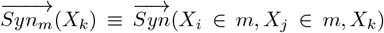. Here, 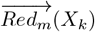 and 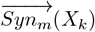 are vector-wise values of HOI *>* 0 and HOI *<* 0, respectively, for the triplet (*X*_*i*_, *X*_*j*_, *X*_*k*_), where *X*_*i*_ and *X*_*j*_ are nodes belonging to the macroregion *m*, and *X*_*k*_ the set of all nodes in the atlas (*k* = 1, …, *N*). Notice that here HOI can indicate either interaction values coming from the O-information or from the gradients.

As a result, for each SC-HOI modular coupling analysis, we obtained 3D redundancy and synergy brain maps that were represented by ℛ^*m*^, defined as 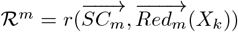, and 𝒮^*m*^, defined as 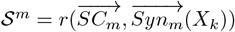 Separated intra-modular and inter-modular contributions to ℛ^*m*^ and 𝒮^*m*^ were also calculated by accounting the non-zero elements within the macroregion mask (intra-modular) and between the macroregion and the rest of the brain (inter-modular).

Multiple comparisons correction was applied in each of the brain maps using FDR correction.

## Results

We separately assessed the structural coupling with redundancy-dominated (ℛ^*m*^) and synergy-dominated (𝒮^*m*^) interactions for different modules *m*, computed using both the O-information and their gradients as separated metrics. A sketch of the complete methodology is given in Figure 1.

**Figure 1.**
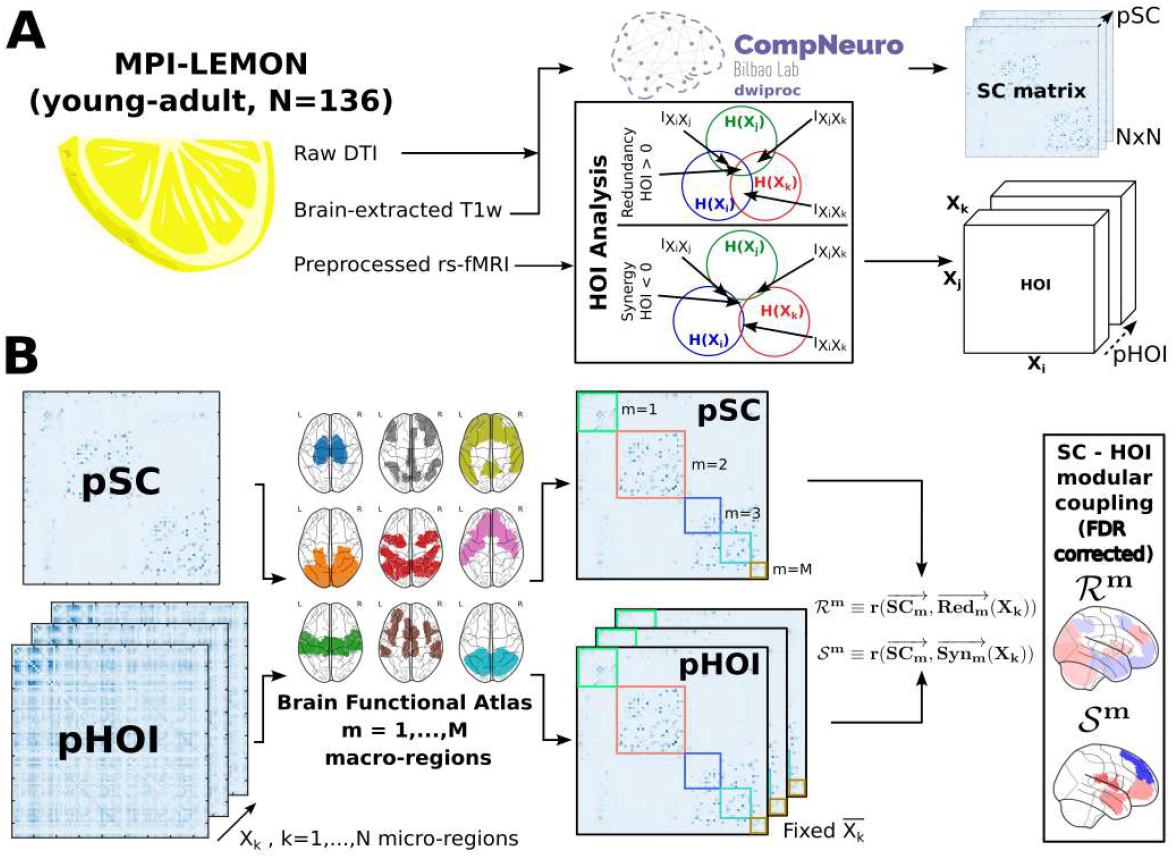
Methodological overview and workflow. **(A)** Time series for each network node (micro-region) were extracted from preprocessed rs-fMRI data, and HOI metrics were computed: O-information and gradients of O-information. For both metrics, positive values indicate a dominance of redundancy, and negative values indicate a dominance of synergy. Population functional HOI matrices were obtained by the median value at each triplet across participants. Population SC matrices were obtained by the median value across participants at a given position. **(B)** Using a functional atlas with *N* micro-regions (nodes in the brain network) and *M* macroregions, that here corresponds to the classical resting state networks, we obtained brain maps of SC-HOI modular coupling, ℛ^*m*^ or 𝒮^*m*^, as the Spearman’s correlation between 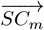 and 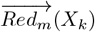 or 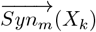. 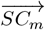 represents the vector-wise structural connectivity between the pairs of micro-regions in the macroregion *m*, while 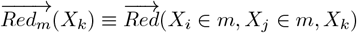, and 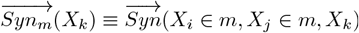, the vector-wise values of HOI*>*0 and HOI*<* 0 for the triplet (*X*_*i*_, *X*_*j*_, *X*_*k*_), where *X*_*i*_ and *X*_*j*_ are a pair of micro-regions belonging to the macroregion *m*, and *X*_*k*_ any micro-region in the atlas.

As a reminder, the results reported below refer to the correlation between SC and the corresponding pair within the module which forms an informational triplet with a third variable, within or outside the module. FDR correction with an alpha level of 0.05 was used to assess the statistical significance of the correlations.

### Nodes and Macroregions defined by an anatomical atlas

#### Structural correlates of O-information

Before evaluating the coupling between SC and functional HOI at the modular level, we first performed a global analysis considering the entire brain as a single macroregion (Figure 2A1). For redundant interactions, we observed significant correlations with SC across nearly all brain regions, with the strongest associations in the right cerebellum (*r* = 0.357, *p* = 10^−16^), left thalamus (*r* = 0.345, *p* = 10^−16^), and left cerebellum (*r* = 0.343, *p* = 10^−16^). Regarding synergistic interactions, the highest positive correlations with SC were identified in the left accumbens (*r* = 0.229, *p* = 10^−16^), right parahippocampal region (*r* = 0.222, *p* = 10^−16^), and left cerebellum (*r* = 0.218, *p* = 10^−16^).

**Figure 2.**
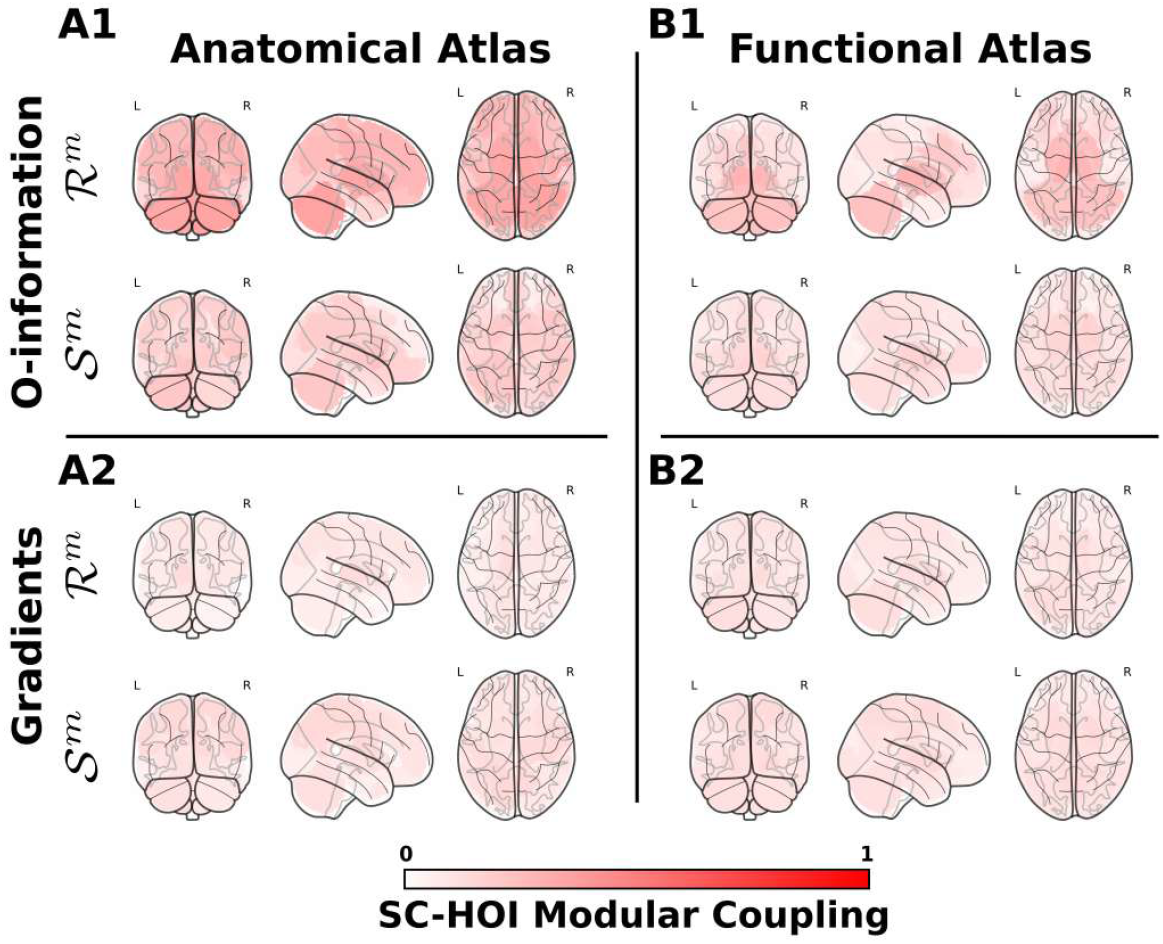
Structural connectivity coupling with High-Order Interactions for the entire brain considered as the only module. **(A1)** Brain maps of modular SC-HOI coupling ℛ^*m*^ and 𝒮^*m*^ based on O-information, obtained by the the Spearman’s correlation between 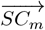 and 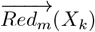, and between 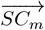 and 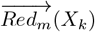, where network nodes are defined by using an anatomical atlas. **(A2)** Brain maps of modular SC-HOI coupling ℛ^*m*^ and 𝒮^*m*^ derived from gradients, obtained by the the Spearman’s correlation between 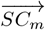 and 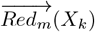, and between 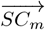 and 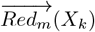 using network nodes from an anatomical atlas. **(B1**,**B2)** Similar to panels A1 and A2, but network nodes are defined from a functional atlas. Therefore, regardless of whether brain regions are defined using an anatomical or functional atlas, when brain modules are not considered in the analyses, the structural coupling associated with both redundancy and synergy consistently exhibits positive correlation values.

Next, we evaluated the coupling between SC and high-order redundant and synergistic interactions at the modular level. The results for anatomical modules are presented in Figure 3A.

**Figure 3.**
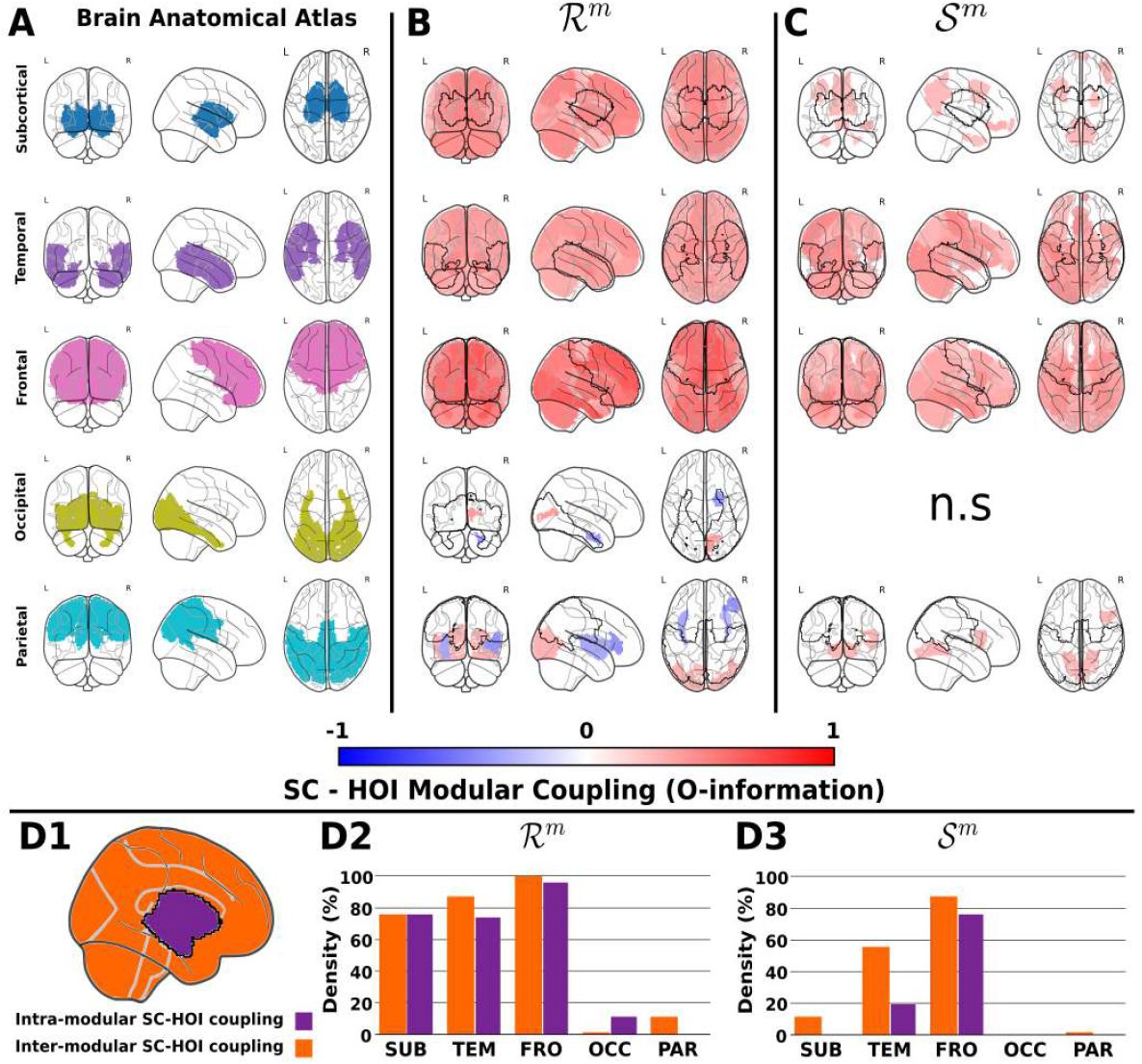
Structure - Function Coupling for different macroregions defined from an anatomical atlas, and high-order interactions defined from the O-information. **A:** Macroregions or modules obtained from network nodes obtained from an anatomical atlas. **B:** Brain maps of modular SC-HOI coupling ℛ^*m*^, obtained by the the Spearman’s correlation between 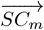 and 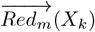 for each one of the defined macroregions. Red color represents positive correlations and blue negative ones. **C:** Brain maps of modular SC-HOI coupling 𝒮 ^*m*^, obtained by the the Spearman’s correlation between 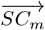 and 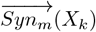 for each one of the defined macroregions. It can be observed that, for the different defined macroregions, an increase in structural connectivity within them may promote either an increase (in red) or a decrease (in blue) in redundancy, corresponding to positive and negative correlations, respectively. In contrast, for synergy, regardless of the macroregion considered, an increase in connectivity within these macroregions is consistently associated with an increase in synergy. **D1:** For visualization purposes of a generic macroregion colored in purple, masks of intra-modular (purple) and inter-modular (orange) contribution. **D2:** Redundant intra- and inter-modular contributions within each macroregion, calculated by the density of non-zero elements of ℛ^*m*^ contained respectively within or outside the intra-modular mask. **D3** Similar than D2 but for 𝒮 ^*m*^. “n.s.” indicates no significant results. All correlation values depicted here are statistically significant after FDR correction (p*<*0.05).

For redundant interactions, we observed a consistent structural correspondence within the sub-cortical macroregion, as well as in the temporal and frontal poles (Figure 3B). Specifically, we found significant structural correlates across most brain regions, with the strongest correlations occurring in the subcortical-thalamus right (*r* = 0.476, *p* = 10^−16^), temporal-thalamus right (*r* = 0.473, *p* = 10^−16^), and frontal-hippocampus left (*r* = 0.47, *p* = 10^−16^) interactions. A distinct pattern emerged in the occipital and parietal regions, where correlations with the structure are both positive and negative. In the occipital lobe, the occipital pole showed a positive correlation with the right pars orbitalis (*r* = 0.39, *p* = 0.043) but a negative correlation with the right corpus callosum (*r* = −0.393, *p* = 0.021). In the parietal lobe, significant correlations were observed: positive with the left isthmus cingulate (*r* = 0.292, *p* = 0.016), and negative with the right pars opercularis (*r* = −0.421, *p* = 10^−3^).

When examining the coupling between structural connectivity (SC) and synergistic interactions (𝒮) (Figure 3C), we observed a similar pattern to that found for redundant interactions, particularly in the temporal and frontal poles. More specifically, SC correlates within these poles were evident across most regions of the brain. The strongest significant correlations were identified between the temporal pole and the left caudal anterior cingulate (*r* = 0.455, *p* = 10^−16^), as well as between the frontal pole and the left middle temporal gyrus (*r* = 0.499, *p* = 10^−16^). In contrast, subcortical and parietal modules exhibited more localized correlation patterns. The highest correlations were found between the subcortical macroregion and the right paracentral lobule (*r* = 0.309, *p* = 0.006), and between the parietal pole and the right parahippocampal gyrus (*r* = 0.278, *p* = 0.039). No significant correlations were found for the occipital pole. As a measure of global module segregation and integration, we computed the intra-modular and inter-modular contributions for both redundant and synergistic SC coupling (Figure 3D1). The densities of redundant and synergistic intra- and inter-modular connections for each macroregion are presented in Figures 3D2-D3, respectively.

Overall, we found that the subcortical, temporal, and frontal macroregions exhibited a balance between intra-modular and inter-modular coupling in the case of redundancy. However, this balance did not hold for synergy, where it was only observed in the frontal macroregion. In contrast, both the temporal and subcortical regions were primarily characterized by inter-modular coupling.

### Structural correlates of Gradients of O-information

As a complementary analysis for the HOI captured by the O-information, we extended our results by assessing the gradients of O-information of order 3. By examining these gradients we can understand how local changes in the system, in this case addition of a triplet of variables, affect the overall information structure.

For the scenario in which the entire brain was considered as a single macroregion, we observed significant correlations across the whole brain for both redundant (ℛ) and synergistic (𝒮) interactions (Figure 2A2). For redundant interactions, the highest correlations were found in the left thalamus (*r* = 0.134, *p* = 10^−7^), left caudate (*r* = 0.1199, *p* = 10^−7^), and left inferior temporal gyrus (*r* = 0.103, *p* = 10^−7^). Regarding synergistic interactions, significant correlations were observed in the left superior frontal gyrus (*r* = 0.15, *p* = 10^−16^), right pars triangularis (*r* = 0.149, *p* = 10^−16^), and right posterior cingulate cortex (*r* = 0.147, *p* = 10^−16^).

When examining the modular coupling between SC and redundancy-derived gradients, we found that the frontal pole exhibited significant correlations across almost the entire brain (Figure 4B). The strongest correlation was observed with the left rostral middle frontal gyrus (*r* = 0.452, *p* = 10^−16^). In contrast, within the subcortical, temporal, occipital, and parietal modules, correlation values were more regionally localized. The highest correlations were found in the following interactions: subcortical-putamen right (*r* = 0.292, *p* = 0.012), temporal-accumbens right (*r* = 0.332, *p* = 0.007), occipital-postcentral right (*r* = 0.425, *p* = 0.041), and parietal-precentral right (*r* = 0.339, *p* = 0.018).

**Figure 4.**
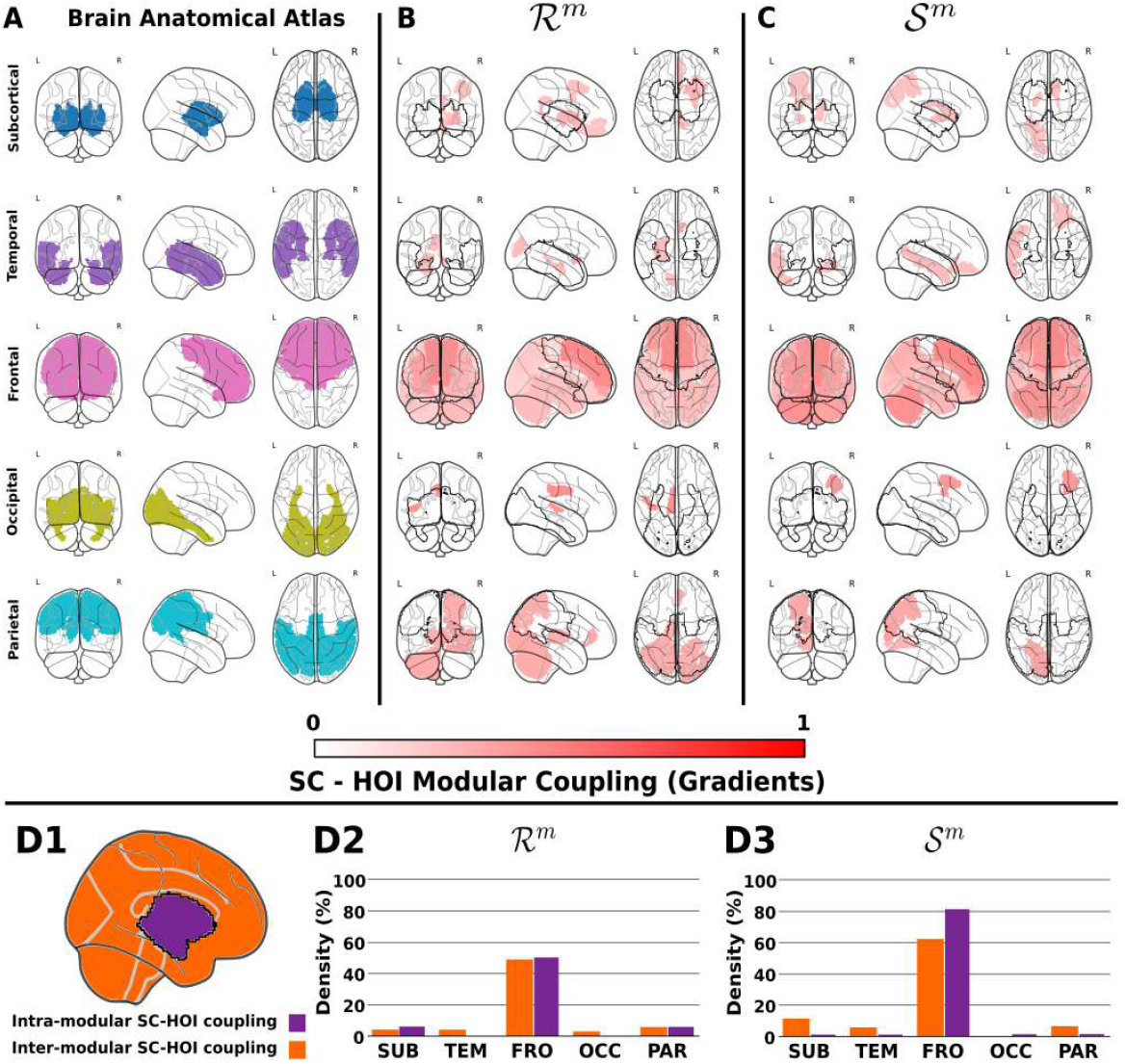
Structure - Function Coupling for different macroregions defined from an anatomical atlas, and high-order interactions defined from the Gradients of O-information. The different panels A, B, C, D1, D2 and D3 represent similar information as the ones in Figure 3, but now R and S are obtained from the gradients of the O-information (for details, see methods). All correlation values depicted here are significant after FDR correction (p*<*0.05).

In relation to synergistic interactions derived from gradients (Figure 4C), the modular coupling between exhibited a similar pattern to that observed for redundant interactions. Specifically, we found that the frontal pole displayed a widespread interaction pattern, with the strongest correlation occurring with the right rostral anterior cingulate (*r* = 0.463, *p* = 10^−16^). For the subcortical, temporal, occipital, and parietal modules, correlation patterns were more regionally localized. The highest correlations were observed in the following interactions: subcortical-pallidum left (*r* = 0.273, *p* = 0.011), temporal-medial orbitofrontal left (*r* = 0.272, *p* = 0.036), occipital-caudal anterior cingulate right (*r* = 0.399, *p* = 0.048), and parietal-corpus callosum (*r* = 0.371, *p* = 0.04).

Finally, we computed the individual intra-modular and inter-modular contributions of gradient-derived redundancy and synergy (Figure 4D2-D3). Interestingly, and in contrast to the results obtained from O-information, the gradient-based interactions revealed that only the frontal macroregion maintained a balance between inter-modular and intra-modular coupling. In contrast, the other macroregions exhibited minimal and statistically insignificant levels of interaction.

### Nodes and Macroregions defined by a functional atlas

#### Structural correlates of O-information

Firstly, as previously done for anatomical macroregions, the first step was to examine the structural correlates of ℛ and 𝒮 without partitioning the brain, considering it as a single region (Figure 2B1).

For redundant interactions, we found significant correlations with SC across the entire brain, with the strongest correlations observed in the left thalamus (*r* = 0.295, *p* = 10^−16^), right thalamus (*r* = 0.282, *p* = 10^−16^), and left putamen (*r* = 0.259, *p* = 10^−16^). For synergistic interactions, only positive correlations with SC were identified, with the highest values found in the right pallidum (*r* = 0.187, *p* = 10^−16^), right putamen (*r* = 0.171, *p* = 10^−16^), and left accumbens (*r* = 0.169, *p* = 10^−16^).

Next, we assessed the modular coupling between SC and redundancy using macroregions defined in the functional atlas (Figure 5A). For redundant interactions (Figure 5B), we found that the subcortical, frontoparietal, and default mode networks (DMN) exhibited a similar pattern of widespread correlation values across the brain. The highest correlation values were observed in the following interactions: subcortical-thalamus right (*r* = 0.535, *p* = 10^−16^), frontoparietal-somatomotor (*r* = 0.574, *p* = 10^−16^), and DMN-caudate (*r* = 0.524, *p* = 10^−16^). In contrast, the visual, somatomotor, dorsal attention, and ventral attention networks showed a more localized correlation pattern. The most prominent interactions were: visual-visual (*r* = 0.331, *p* = 0.002), somatomotor-ventral diencephalon (*r* = 0.356, *p* = 0.003), dorsal attention-DMN (*r* = 0.412, *p* = 10^−4^), and ventral attention-frontoparietal (*r* = 0.526, *p* = 10^−4^). The limbic network was not involved in any significant correlation.

**Figure 5.**
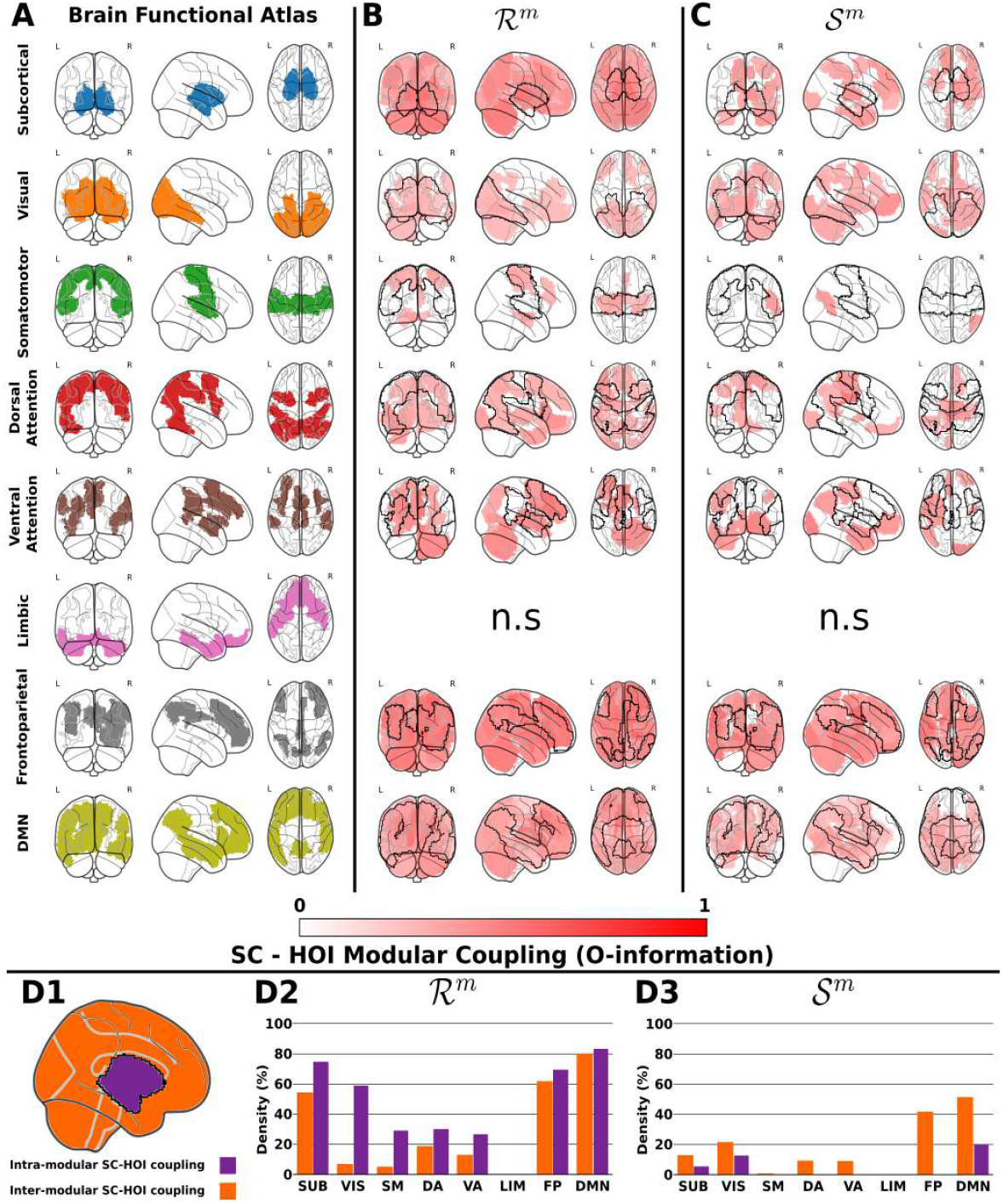
Structure - Function Coupling for different macroregions defined from a functional atlas, and high-order interactions defined from the O-information. **A:** Functional macroregions corresponding with classical Yeo’s Resting State Networks and one additional subcortical network, obtained from a composite of several subcortical regions obtained from the Freesurfer segmentation. **B:** Brain maps of modular SC-HOI coupling ℛ^*m*^, obtained by the the Spearman’s correlation between 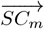 and 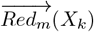 for each one of the defined macroregions. Red color represents positive correlations and blue, negative ones. **C:** Brain maps of modular SC-HOI coupling 𝒮 ^*m*^, obtained by the the Spearman’s correlation between 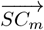 and 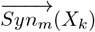 for each one of the defined macroregions. **D1-D3:** Similar to panels in Figure 3, but now the masks for intra-modular and inter-modular contributions correspond to functional macroregions. All correlation values depicted here are significant after FDR correction (p*<*0.05).

For functional modules, the coupling between SC and synergy exhibited a widespread pattern of significant correlations throughout the brain (Figure 5C). The highest correlation values were observed in the following interactions: frontoparietal-somatomotor (*r* = 0.569, *p* = 10^−4^) and DMN-visual (*r* = 0.384, *p* = 10^−16^). In contrast, the remaining networks displayed a more localized correlation pattern. The strongest correlations were found in the following interactions: subcortical-thalamus (*r* = 0.411, *p* = 0.019), visual-DMN (*r* = 0.408, *p* = 10^−4^), somatomotor-dorsal attention (*r* = 0.372, *p* = 0.013), dorsal attention-frontoparietal (*r* = 0.503, *p* = 10^−4^), and ventral attention-visual (*r* = 0.476, *p* = 0.25).

For redundant and synergistic SC coupling, the individual intra-modular and inter-modular contributions are depicted in Figure 5D1. The profiles of redundant and synergistic intra- and inter-modular interactions for each functional macroregion are shown in Figures 5D2-D3, respectively. Notably, for redundancy (ℛ), we observed a balance between intra-modular and inter-modular interactions in the subcortical, frontoparietal, and DMN networks. In contrast, for synergy (𝒮), no network exhibited such a balance. Remarkably, the DMN displayed a higher degree of inter-modular coupling compared to intra-modular interactions.

#### Structural correlates of Gradients of O-information

When no functional modules were used and the entire brain was considered as a single macroregion, a similar pattern of significant correlations was observed for both redundancy and synergy derived from gradients of O-information (Figure 2B2). For redundancy, the highest correlations were found in the caudate (*r* = 0.139, *p* = 10^−16^), cerebellum (*r* = 0.128, *p* = 10^−16^), and ventral attention network (*r* = 0.12, *p* = 10^−16^). For synergistic interactions, positive correlations with SC were identified in regions such as the putamen (*r* = 0.149, *p* = 10^−16^) and the dorsal attention network (*r* = 0.145, *p* = 10^−16^).

When defining multiple functional macroregions, as shown in Figure 6A, we first assessed the coupling between SC and redundancy (Figure 6B). We found a similar pattern of significant correlations in the dorsal attention, frontoparietal, and default mode networks (DMN), with the highest values observed in the following interactions: dorsal attention-DMN (*r* = 0.449, *p* = 10^−4^), frontoparietal-ventral attention (*r* = 0.536, *p* = 10^−14^), and DMN-limbic (*r* = 0.334, *p* = 10^−4^). In contrast, the visual, somatomotor, and ventral attention networks exhibited a more localized correlation pattern. The strongest correlations were found in: visual-visual (*r* = 0.288, *p* = 0.016), somatomotor-somatomotor (*r* = 0.381, *p* = 0.009), and ventral attention-somatomotor (*r* = 0.415, *p* = 0.009). The subcortical and limbic networks did not show any significant correlations.

**Figure 6.**
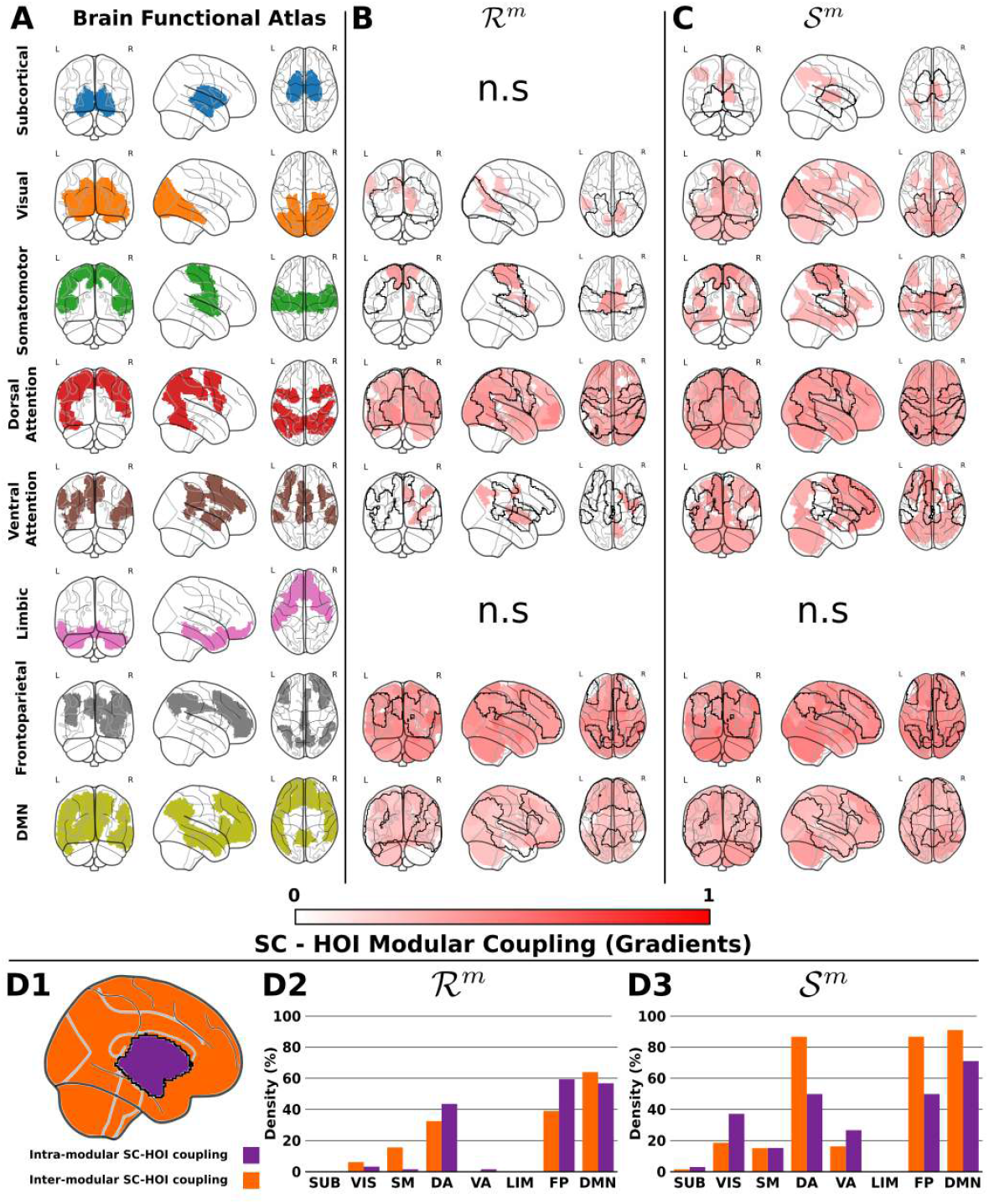
Structure - Function Coupling for different macroregions defined from a functional atlas, and high-order interactions defined from the Gradients of O-information. The different panels here A, B, C, D1, D2 and D3 show similar results as shown in Figure 4 but now R and S are estimated from the gradients. All correlation values depicted here are significant after FDR correction (p*<*0.05).

Regarding synergistic interactions (Figure 6C), the dorsal attention, ventral attention, frontoparietal, and default mode networks (DMN) exhibited a widespread pattern of significant correlations. The strongest interactions were observed in: dorsal attention-DMN (*r* = 0.467, *p* = 10^−4^), ventral attention-DMN (*r* = 0.459, *p* = 0.002), frontoparietal-accumbens (*r* = 0.589, *p* = 10^−16^), and DMN-cerebellum (*r* = 0.374, *p* = 10^−16^). In contrast, the subcortical, visual, and somatomotor networks displayed a more localized correlation pattern. The highest correlations were found in: subcortical-thalamus (*r* = 0.317, *p* = 0.007), visual-visual (*r* = 0.367, *p* = 10^−4^), and somatomotor-somatomotor (*r* = 0.422, *p* = 0.002).

For the coupling between redundancy and synergy derived from gradients of O-information, the individual intra-modular and inter-modular contributions are depicted in Figure 6D1. The densities of redundant and synergistic intra- and inter-modular interactions for each macroregion are shown in Figures 6D2-D3, respectively. Overall, for redundancy, we observed a balance between intra-modular and inter-modular coupling in the dorsal attention, frontoparietal, and default mode networks (DMN). However, this balance was not present for synergy, as no network exhibited a similar equilibrium between intra-modular and inter-modular couplings. Notably, for the dorsal attention, frontoparietal, and DMN networks, inter-modular contributions—representing interactions with the rest of the brain—were greater than intra-modular contributions, which reflect interactions occurring within each network.

## Discussion

This study investigated how the structural connectivity (SC) is related to functional high-order interactions (HOI) in the human brain. Our results show that while redundancy can increase between nodes with strengthened SC (confirming previous findings (Andrea I. Luppi, P. A. M. Mediano, et al. 2022)), it can also decrease, displaying an overall inconsistent relationship. In contrast, results revealed that synergy consistently increases with stronger SC. Crucially, these results were only observed when examining the coupling between SC and functional HOI at a modular level, as these interactions did not emerged in our analyses unless the brain was explicitly modularized. These findings were verified to be independent of brain parcellation, and to be robust across different definitions of brain modules. Previous research on the relationship between SC and FC has been primarily focused on pairwise interactions, leaving the coupling between SC and functional HOI beyond pairwise relationships largely unexplored. On the other hand, recent studies (Andrea I. Luppi, P. A. M. Mediano, et al. 2022; Varley, Pope, Faskowitz, et al. 2023; Varley, Pope, Maria Grazia, et al. 2023; Scagliarini, Nuzzi, et al. 2023; Scagliarini, Sparacino, et al. 2024) have utilized HOI metrics such as partial entropy decomposition, and gradients of O-information to identify redundant and synergistic subsets within brain networks. These analyses are enabling deeper insights into the interplay between structural and functional connectivity that cannot be captured through pairwise interactions alone. In this study, we contribute to this line of work by investigating the relationship between SC and functional HOI, aiming to elucidate how structural connectivity correlates with high-order functional interactions.

Previous work has shown that both redundancy and synergy tends to correlate with SC, with redundancy exhibiting a stronger correlation (Andrea I. Luppi, P. A. M. Mediano, et al. 2022). The primary distinction between this work and ours lies in the way that HOI is computed: Luppi et al. employed integrated information decomposition (P. A. M. Mediano et al. 2021) to measure information flow from the past to the future within a system, thus assessing HOI in the temporal domain. In contrast, our approach calculates HOI across spatial interactions involving triplets of variables using O-information and its gradients, but not across time. This methodology was chosen to focus on the structural-functional coupling between SC and redundancy and synergy in a purely spatial aspect.

Our findings demonstrate that both redundancy-dominated (ℛ) and synergy-dominated (𝒮) interactions exhibit significant correlations with SC. These correlations can be both positive and negative for ℛ, whereas for 𝒮, they are exclusively positive. Thus, in addition to the findings reported in (Andrea I. Luppi, P. A. M. Mediano, et al. 2022), which showed that SC drives redundant interactions, we reveal that SC can also be associate with synergistic interactions. Another important characteristic of our approach is the focus on modular SC-HOI coupling. This distinction is clearly demonstrated by the fact that, when no brain partitioning into modules was applied and functional HOI was computed for the entire brain, ℛ never exhibited negative correlations with SC, consistent with (Andrea I. Luppi, P. A. M. Mediano, et al. 2022). In contrast, our results, where SC correlates both positively and negatively with ℛ, emerge as a direct consequence of assessing the modular SC-HOI correspondence. From a methodological perspective, we remark that this study assesses statistical similarities and dependence between SC and functional HOI, and thus the term *coupling* does not imply an underlying mechanism or causal relationship (Liu et al. 2022).

When analyzing redundancy-dominated functional HOI calculated via O-information using the anatomical atlas, we observed a consistent pattern in the subcortical, temporal, and frontal modules, where redundant interactions positively correlated with SC—a trend that extended across the entire brain. In contrast, the occipital and parietal modules exhibited a different pattern, characterized by more localized correlations and negative associations with SC. This suggests that regions with higher redundant interactions might be less related to structure. Regarding synergistic functional HOI, the temporal and frontal modules displayed a pattern similar to that observed for redundant interactions, with positive correlation values widely distributed across the brain. Meanwhile, the subcortical and parietal modules exhibited more regionally specific correlation patterns.

Using the functional atlas, we observed that the subcortical, visual, somatomotor, dorsal attention, ventral attention, and frontoparietal networks exhibited higher values of intra-modular SC-HOI coupling. This suggests that these networks play a segregating role in redundant interactions. The default mode network (DMN) displayed a predominant redundant role, functioning both in integration and segregation, consistent with previous findings across the lifespan (Camino-Pontes et al. 2018). Conversely, when analyzing synergistic interactions, all networks contributed more prominently to integration in brain interactions.

The consistency of our results, demonstrating that both ℛ and 𝒮 can exhibit significant coupling with SC, was independent of the chosen brain partition. Thus, whether studying HOI coupling through O-information or its gradients, and regardless of whether anatomical or functional brain partitions were used, these findings remained consistent. Furthermore, the diverse repertoire of distinct patterns of negative and positive correlations with SC, depending on the modular organization of functional HOI, represents one of the key contributions of this work. A promising future direction would be to investigate how these patterns may vary across different brain pathologies. Additionally, exploring alternative approaches to quantify synergy and redundancy presents another valuable extension of this research. For instance, methodologies such as information flow analysis (Andrea I. Luppi, P. A. M. Mediano, et al. 2022) or time-resolved approaches (Varley, Pope, Maria Grazia, et al. 2023; Pope et al. 2024) could be employed. Moreover, investigating higher-order interactions beyond three-way interactions— which have been shown to be critical in other contexts—represents another compelling avenue for future research (Herzog et al. 2022).

Overall, this work represents a first exploration into the relevance of coupling of SC with functional relationships that go beyond pairwise interactions. Our results revealed that the association between SC and functional HOI is more complex than previously thought, having a consistent relationship with synergy but an inconsistent one with redundancy. We hope that our findings will encourage further research in this promising area of investigation.

## Acknowledgments

BCP was supported by a grant from the Basque Government (PRE-20202-1-0187). IFI acknowledges financial support from a predoctoral grant from the Basque Government (PRE-2024-2-0208). AE is supported by the Spanish Ministry of Science and Innovation grant (RYC2021-032390-I). JMC and AE are founded by the Spanish Ministry of Science (grant PID2023-1480120B-I00). JMC, PB, ID and AE are founded by Ikerbasque: The Basque Foundation for Science. JMC is founded by the Department of Economic Development and Infrastructure of the Basque Country (Elkartek Program, grant KK-2021-00009) and the Health Department of the Basque Country (grants 2022111031 and 2023111002).

